# Custom-engineered hydrogels for delivery of human iPSC-derived neurons into the injured cervical spinal cord

**DOI:** 10.1101/2023.05.10.540225

**Authors:** V.M. Doulames, L.M. Marquardt, M.E. Hefferon, N. Baugh, R.A. Suhar, A.T. Wang, K. Dubbin, J.M. Weimann, T.D. Palmer, G.W. Plant, S.C. Heilshorn

## Abstract

Cervical damage is the most prevalent type of spinal cord injury clinically, although few preclinical research studies focus on this anatomical region of injury. Here we present a combinatorial therapy composed of a custom-engineered, injectable hydrogel and human induced pluripotent stem cell (iPSC)-derived deep cortical neurons. The biomimetic hydrogel has a modular design that includes a protein-engineered component to allow customization of the cell-adhesive peptide sequence and a synthetic polymer component to allow customization of the gel mechanical properties. *In vitro* studies with encapsulated iPSC-neurons were used to select a bespoke hydrogel formulation that maintains cell viability and promotes neurite extension. Following injection into the injured cervical spinal cord in a rat contusion model, the hydrogel biodegraded over six weeks without causing any adverse reaction. Compared to cell delivery using saline, the hydrogel significantly improved the reproducibility of cell transplantation and integration into the host tissue. Across three metrics of animal behavior, this combinatorial therapy significantly improved sensorimotor function by six weeks post transplantation. Taken together, these findings demonstrate that design of a combinatorial therapy that includes a gel customized for a specific fate-restricted cell type can induce regeneration in the injured cervical spinal cord.

## Introduction

Cervical spinal cord injury (SCI) results in a devastating and permanent loss of sensorimotor function and, to date, there is no cure. Here, we describe a combinatorial therapy specifically designed to treat the sensorimotor dysfunction that defines cervical SCI: the intraspinal transplantation of human stem cell-derived deep cortical neurons delivered in a biomimetic, injectable hydrogel.

There are key regional anatomical and functional differences along the length of the spinal cord, and each region warrants careful consideration of the therapeutic goals and design criteria to meet these goals.^1^ Only 12% of preclinical SCI research is focused on cervical SCI, although over half of all SCIs occur within the cervical region.^2, 3^ This is largely because the cervical spinal cord is more prone to injury than lower spinal segments. The cervical spine is more mobile than the lower spine, but is encased by smaller vertebrae and lacks external stabilization such as the perivertebral muscle and ribcage, which provides protection to the thoracic spine.^4–6^ Importantly, improvements in emergency medicine have dramatically increased survival in patients with cervical SCI making this a rapidly growing demographic, but therapies to improve long-term functional outcomes have not progressed in parallel. Lower cervical SCIs are associated with worse prognosis; patients with lower cervical SCIs are more likely to receive a Grade A score (complete impairment) within the American Spinal Injury Association Impairment Scale and tend to have a lower median motor score at presentation.^4^ Designing therapies to improve function and quality of life are necessary.

Intraspinal cell transplantation therapies have emerged as a promising and customizable way to treat primary and secondary injury cascades inherent to cervical SCI; however, these therapies are logistically challenging, which reduces efficacy and translational potential.^7–10^ First, there is no clear consensus in the field as to what the optimal transplanted cell type is to promote recovery post SCI. In stem cell-derived therapies, transplanting developmentally immature progenitor cells may lead to better cell survival during dissociation and transplantation, but they may not differentiate into a desirable cell type *in situ*, which can limit therapeutic effectiveness.^11^ On the other hand, transplantation of mature cell types offers more control over final lineage commitment, but the cells are more sensitive to the manipulations required prior to transplantation.

Second, almost all cell transplantation paradigms are traditionally plagued by low viability, with reported survival rates as low as 1%.^11, 12^ Cell injection via syringe minimizes iatrogenic injury (*i.e.* injury caused by more invasive surgical protocols), but this method subjects the transplanted cells to shear forces, resulting in mechanical membrane damage, which significantly contributes to cell death. We and others have shown that cells injected in an aqueous fluid solution, such as saline, are particularly susceptible to both shear and extensional forces within the syringe.^13–18^ In addition to membrane damage and cell death during acute injection, reflux along the injection needle tract is an additional obstacle to delivering cells to their intended target. Due to positive pressure within the spinal cord, transplanted cells reflux down the path of least resistance (*i.e.* up the needle tract), leading to fewer cells deposited at the target site.^19^ Finally, the transplanted cells must contend with the lack of extracellular matrix within the injury site. Following SCI, an irregular-shaped cystic cavity can form, and transplanted cells need to fill this void space, which requires a physical scaffold to bridge the lesion and integrate with the native tissue.^20^

Using engineered hydrogels to both encapsulate and deliver cells can improve the efficacy of cell transplantation through biochemical and biomechanical interactions.^21^ To address cell damage during acute injection, injectable hydrogels have been created to protect cells from shear-induced membrane damage. These systems function by enabling the bulk of the hydrogel to move through the needle as a solid plug, with only the edges adjacent to the needle wall flowing like a liquid, limiting shear-stress exposure to a very small fraction of the total volume. As a result, the majority of cells pass through the needle without experiencing cell membrane damage.^17^ To address reflux and dispersion away from the injection site, injectable hydrogels have been designed to have rapid shear-thinning and self-healing behavior.^22^ This combination of properties allows for smooth injection and quick reformation into a gel in situ to keep cells at the injection site.^14^ To promote cell viability within the injured tissue, hydrogels can be engineered to provide biochemical cues that mimic the native extracellular matrix and present ligands that promote cell migration and neurite extension into the host tissue.^17^ These parameters can all be manipulated to best suit the injection protocol, target injection site, and transplanted cell type.

Several cell types with differing intended therapeutic function have been explored as cell transplantation therapies for SCI. Tailoring the delivery vehicle to the biological properties of the cell type can improve transplanted cell survival and functional integration.^23–26^ We previously developed and optimized a modular hydrogel delivery system (SHIELD; shear-thinning hydrogel for injectable encapsulation and long term delivery) that improved transplantation of Schwann cells for SCI.^14^ In preclinical and clinical studies, transplanted Schwann cells aid in halting the secondary injury cascade and preventing further tissue damage.^27–33^ Consistent with this intended therapeutic function, we found that Schwann cells delivered in an optimized SHIELD formulation resulted in statistically significant decreases in cystic cavity volume compared to Schwann cells delivered in saline in a rodent model of SCI.

The advancement of iPSC technology has presented a valuable tool for producing differentiated somatic cells with specific characteristics, and numerous investigations exploring the potential applications of cells derived from human iPSCs (hiPSCs) have been launched in recent years.^19^ One line of thought is that replacing lost neurons will allow for repair of severed neurocircuits required for sensorimotor function. There is an enormous amount of cellular diversity within the central nervous system with unique transcriptional signatures that orchestrate the precise functions and neural circuits that shape human behavior.^34^ The long tract connections of cortical projection neurons are an inherent component of the cervical spinal niche, and we hypothesized that this cell type may be especially well suited to integrating into the local neurocircuitry and improving function.^35^ hiPSC-derived deep cortical neurons (hiPSC-DCNs) therefore represent a potential patient-specific regenerative cell type for application within the injured central nervous system (CNS). However, fate-restricted, post-mitotic neurons with long axonal processes are especially mechanosensitive and cannot proliferate – thus, designing methods to promote transplanted cell survival is paramount to determining if this class of cells can integrate and improve function within the adult injured cervical spine. Here we present the customization of an engineered hydrogel delivery vehicle for transplantation of cortical projection neurons and demonstrate its potential to significantly improve the functional outcomes of cell-based therapy in a preclinical model of cervical spinal cord injury.

## Results and Discussion

### The SHIELD family of hydrogels is designed to address the most common delivery problems in cell transplantation therapies

We have designed SHIELD with three specific features to address the key challenges associated with transplanting cells that are particularly sensitive to mechanical and biological stressors: (1) thixotropy to protect the cell membrane during injection, (2) rapid self-healing and *in situ* stiffening to keep cells in place within the injury site, and (3) the inclusion of cell-adhesive ligands to promote cell attachment and functional integration. Our previous work has shown that these individual components of SHIELD can be tuned to both the needs of the transplanted cell type and the therapeutic application.^14, 36, 37^

SHIELD is a two-component material that leverages two stages of cross-linking (Fig. 1A). The first component is C7, an engineered recombinant protein with alternating flexible spacer domains and folded WW domains that serve as crosslinking sites.^38^ We designed three different extracellular matrix-derived ligands into the spacer domain of the C7 protein with the goal of identifying the C7 variant that would best support hiPSC-DCN survival (full-length sequences provided in **Table. S1**). The second component is a multi-arm polyethylene glycol (PEG) conjugated with proline-rich peptides (termed a P domain) that physically crosslinks with the WW domains of C7 through peptide-peptide assembly. The PEG-P1 copolymer can be further conjugated with the thermo-sensitive polymer poly(N-isopropylacrylamide) (PNIPAM). At physiological temperature, PNIPAM undergoes hydrophobic collapse to provide secondary physical cross-linking to stiffen the hydrogel network.^36^ By adjusting the concentration of PNIPAM within the SHIELD formulation (0-2.5 wt%), we created a family of hydrogels that span a range of stiffness (*G’* ∼ 10-600 Pa, Fig. 1B) with the goal of selecting the variant that best supports hiPSC-DCN neurite extension.

**Figure 1.**
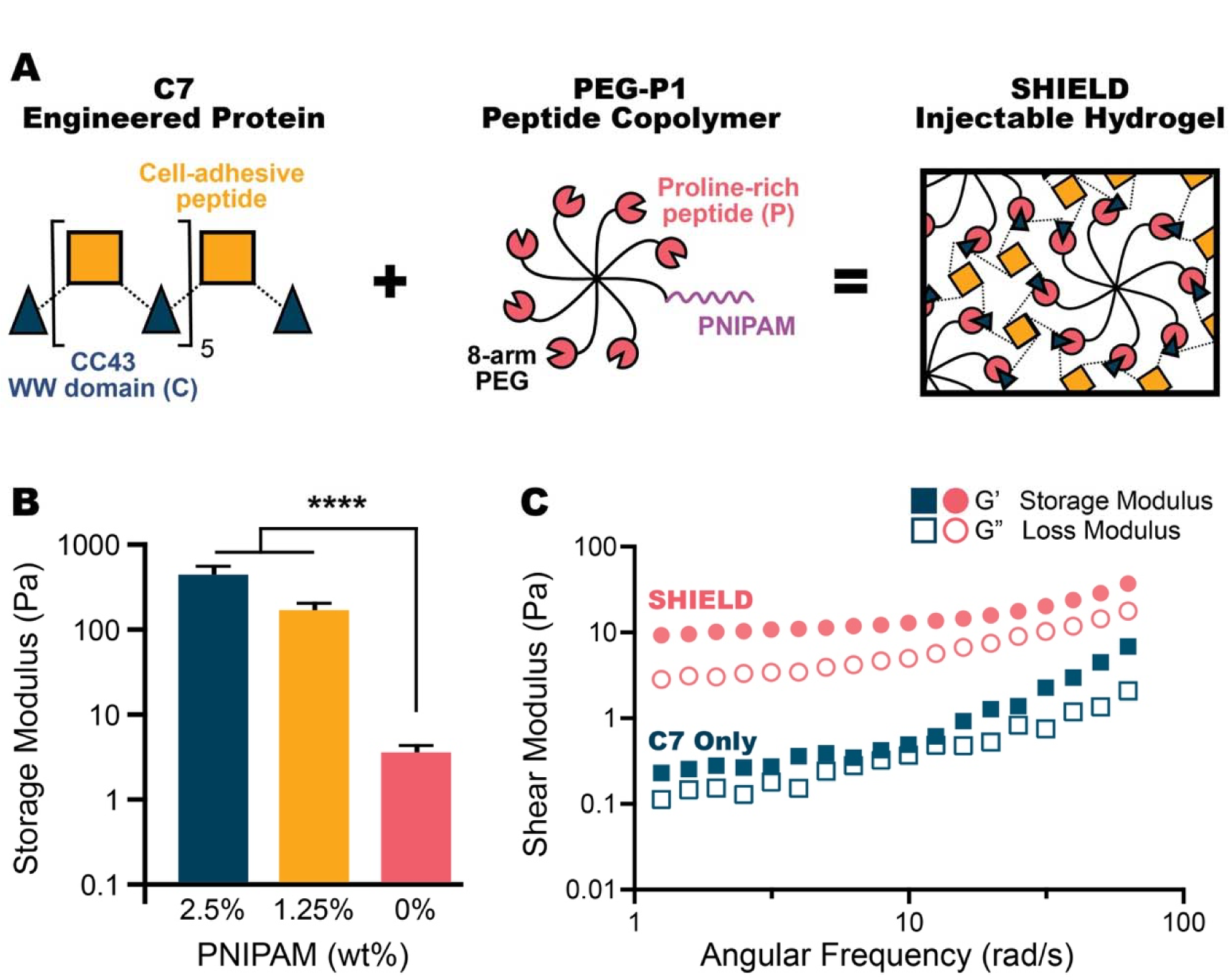
The SHIELD family of hydrogels is designed to address cell loss during transplantation and provide bioactive cues that promote cell survival and integration. **A.** Schematic of the SHIELD system, which is composed of a C7 engineered protein and a PEG-P1 peptide copolymer with varying amounts of PNIPAM copolymer. **B.** The shear storage modulus of SHIELD formulations at 37°C is controlled by tuning the percentage of thermosensitive PNIPAM. Data are mean ± SEM. \*\*\*\**P* < 0.0001; Tukey post hoc test; *n* = 3 to 8. **C.** Representative frequency sweeps of storage (G’) and loss (G”) moduli were performed at 24°C to characterize the viscoelastic behavior of SHIELD (0 wt% PNIPAM) and a negative control sample that includes only the C7 engineered protein without the PEG-P1 binding partner.

These two components engage in two stages of cross-linking. In the first stage *ex situ*, the two components are mixed together in the presence of cells, allowing the two engineered polymers to assemble via reversible, heterodimeric binding of their peptide domains to form a weak gel (Fig. 1C, **Fig. S2**). As a negative control comparison, the measured stiffness of the C7 protein alone is more than an order of magnitude lower, demonstrating that the peptide-peptide interactions are a critical component of the gel mechanical properties (Fig. 1C, **Fig. S2**). When the gel is subjected to a force, the peptide-peptide physical crosslinks are disrupted, allowing the material to shear-thin and flow. After the flow-inducing force is removed, the peptide-peptide bonds rapidly reform, allowing the gel to quickly self-heal. Once injected, the gel warms to body temperature, and the second stage of cross-linking then occurs as the temperature-sensitive PNIPAM undergoes hydrophobic collapse.

### Human iPSCs can be driven towards a cortical neuron phenotype

*In vitro*, human induced pluripotent stem cells (hiPSCs) can undergo directed differentiation that recapitulates the temporal expression of fate-specifying transcription factors during human cortical neurogenesis through an adaptation of the dual SMAD inhibition protocol (Fig. 2A).^35, 39^

**Figure 2.**
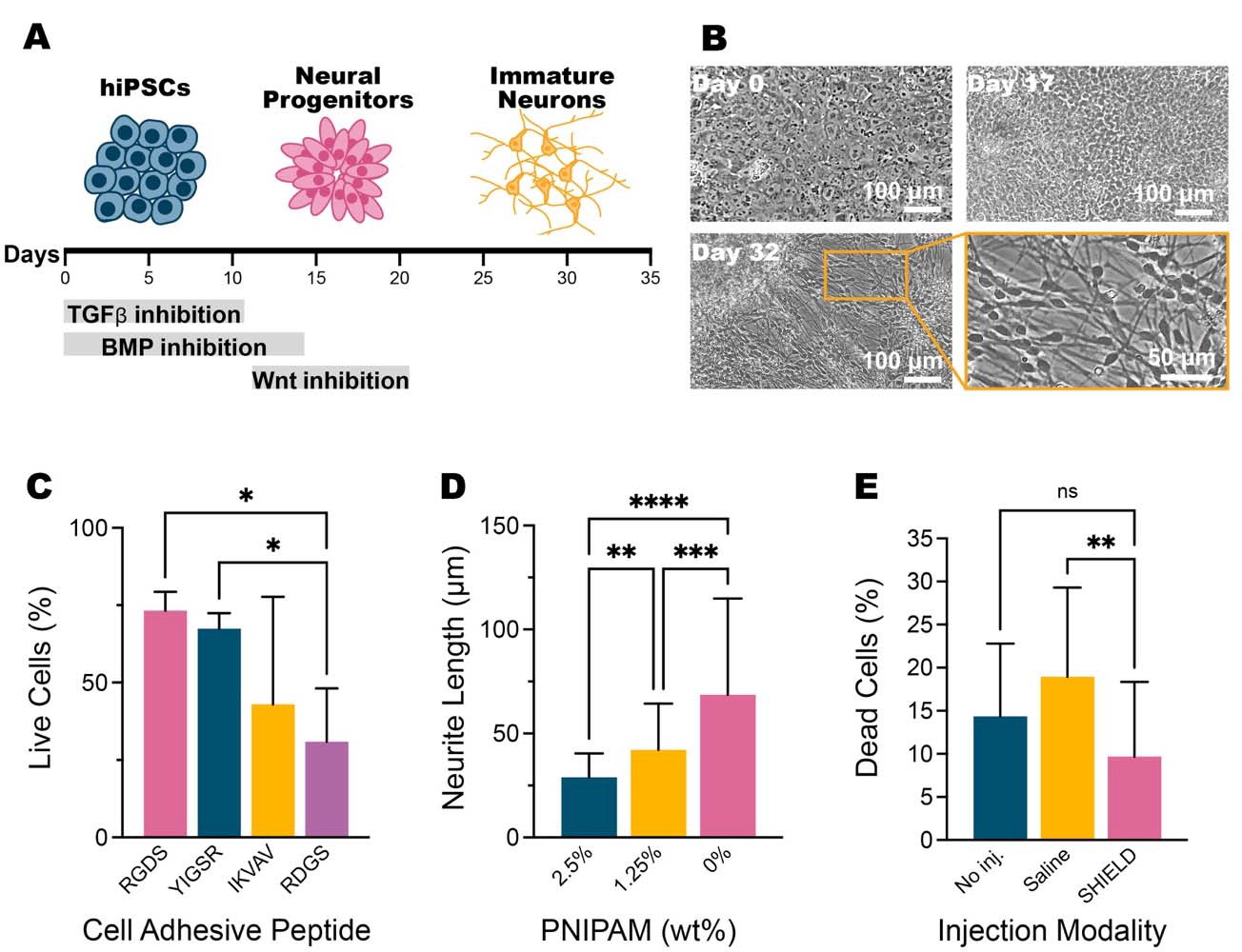
*SHIELD can be customized to promote viability and neurite extension of hiPSC-DCNs* in vitro. **A.** Schematic of the directed differentiation protocol to drive hiPSCs toward a deep cortical neuron (DCN) commitment. **B.** Representative brightfield images of hiPSC cultures at different points of the directed differentiation protocol. At day 0, the culture has a “cobblestone” morphology, consistent with pluripotent stem cells. At day 17, the culture has begun to radially organize into “rosette” structures, consistent with neuroepithelium. By day 32, the culture is largely interconnected by long processes, consistent with maturing neurons. **C.** Percentage of live hiPSC-DCNs (Calcein-AM^+^) after 7 days following encapsulation in SHIELD variants with fibronectin-(RGDS) or laminin-(YIGSR, IKVAV) derived binding domains, or a scrambled non-adhesive control sequence (RDGS). Data are mean ± SD. *P = 0.0150 (RGDS vs. RDGS), *P = 0.0306 (YIGSR vs. RDGS), F = 5.466; one-way analysis of variance (ANOVA) with Dunnet’s multiple comparisons post hoc test; *N* = 2 independent experiments, each with _≥_2 technical replicates. **D.** Quantification of hiPSC-DCN neurite length in SHIELD gels with varying amount of PNIPAM after 3 days in culture. Data are mean ± SD. **P = 0.0020, ***P = 0.0006, ****P < 0.0001, F = 26.40; one-way ANOVA with Dunnet’s multiple comparisons post hoc test; *N* = 2 independent experiments, each with _≥_ 2 technical replicates. **E.** Percentage of dead hiPSC-DCNs (ethidium homodimer-1^+^) following exposure to pipetting in saline (i.e. no injection), injection through a syringe needle (33-G) in saline at 500 nL/min, or injection through a syringe needle (33-G) in SHIELD (RGD binding domain and 0% PNIPAM) at 500 nL/min. Data are mean ± SD. **P = 0.0011, F = 7.648; one-way ANOVA with Dunnet’s multiple comparisons post hoc test; *N* = 2 independent experiments, each with _≥_2 technical replicates.

At day 0, undifferentiated hiPSC cultures demonstrate the morphology of pluripotent stem cells: cuboidal-shaped, tightly packed cells in a cobblestone colony pattern, high nucleus to cytoplasm ratio, and prominent nucleoli (Fig. 2B). To induce differentiation towards a neural fate, hiPSCs are switched to a medium that stimulates neuroectodermal specification and then neurogenesis, followed at later times by the production of glia. The early patterning to neuroectoderm involves the removal of basic fibroblast growth factor (FGF2) and the addition of small molecule inhibitors applied at key timeframes (Fig. 2A), including the inhibition of transforming growth factor beta (TGFb) and bone morphogenetic protein (BMP) signaling.^40, 41^ Within 10 days of neural induction, the hiPSC cultures begin to radially organize into rosette structures, which is a characteristic attribute of neuroepithelial development of the neural tube (Fig. 2B).^42^ *In vivo*, the neuroepithelium of the neural tube is responsible for generating all the neurons and glial cells in the central nervous system during development. Wnt signaling plays a role in correctly patterning neural structures in the dorsal-ventral axis, with high Wnt activity leading to ventralization. To shift the culture towards a dorsal telencephalic identity, porcupine (PORCN) activity was inhibited until day 19, which prevents both canonical and noncanonical Wnt-mediated signaling.^43^

Cultures are maintained for 35 days *in vitro*, at which time most cells have consolidated an anterior cortical fate. Neurogenesis during this period produces predominantly excitatory projection neurons with extensive neurite networks. These immature neurons show robust expression of Type 3 beta tubulin (a microtubule element found almost exclusively in neurons) and MAP2 (a cytoskeletal protein that is abundant in neuronal dendrites) (Fig. 2B, **Fig. S2A**). As seen *in vivo*, neurogenesis precedes gliogenesis, and the vast majority of cells produced during the 35 days of differentiation are young neurons (approximately 85% of the culture is beta tubulin^+^, 15% is Nestin^+^ (marker of neural progenitors), and 0% is positive for glial fibrillary acidic protein (GFAP; marker of glia)) (**Fig. S2B**). If allowed to differentiate for 155 days or longer, glia begin to appear, with approximately 10% of the cells GFAP+, 75% beta tubulin^+^, and 15% Nestin^+^ (**Fig. S2B**). Collectively, this directed differentiation protocol results in a strong enrichment for neurons, and our transplantation strategy utilizes a nearly pure population of cortical neurons and neural progenitors from day 35 of differentiation.

### Tuning biochemical and mechanical cues improves hiPSC-DCN viability and neurite outgrowth

The long neurites of developing neurons (such as hiPSC-DCNs) are particularly sensitive to mechanical strain and applied shear forces during injection. These forces damage the delicate neuroarchitecture and can cause diffuse injuries including stretching or rupture of neurites and damage to the cell body.^44, 45^ Ultimately, this leads to axon degeneration and reduced cell viability.^46^ By day 35 (transplantation age), the majority of hiPSC-DCNs are post-mitotic. Thus, it is critical that cells survive the injection protocol, because these neurons cannot proliferate to replace cells that may be lost during injection.

*In vitro*, survival of these differentiating cells requires cell adhesion to prevent anoikis (*i.e.* programmed cell death in the absence of a matrix), and we postulated that an absence of survival-promoting, matrix-adhesion cues following transplantation may contribute to the low survival of injected cell therapies. We encapsulated hiPSC-DCNs in SHIELD hydrogels formulated from three different C7 variants that included three different cell-adhesive matrix ligands. Specifically, the RGDS peptide sequence derived from fibronectin and the IKVAV and YIGSR peptide sequences derived from laminin were explored, as these epitopes are implicated in a range of cortical neuron behaviors.^47–50^ As a negative control, we also encapsulated cells in a C7 variant that included a non-cell-adhesive, sequence-scrambled RDGS peptide. Over seven days in culture, hiPSC-DCNs grown in SHIELD with the RGDS ligand demonstrated the highest cell viability (Fig. 2C) and the most robust neurite outgrowth (**Fig. S3A**). Therefore, we chose to move forward with SHIELD formulation presenting the fibronectin-derived RGDS binding domain for further studies.

*In vivo*, mechanical signals are important regulators of axon pathfinding for neurons.^5130^ Published studies suggest that reducing the stiffness of a gel increases the speed of neurite outgrowth and branching.^52–54^ By adjusting the percentage of PNIPAM attached to PEG-P1, we created a family of SHIELD hydrogels with a range of stiffness (Fig. 1B). hiPSC-DCNs were cultured in these SHIELD variants of increasing stiffness for three days (all gels included similar concentrations of the RGDS cell-adhesive domain). Beta tubulin^+^ staining demonstrated highest hiPSC-DCN outgrowth in the most compliant gel within the SHIELD family with 0 wt% PNIPAM (Fig. 2D, **Fig. S3B**). Therefore, we chose to move forward with the softest SHIELD formulation for cell transplantation studies.

In recent years, several studies have demonstrated that cell damage during injection protocols that use aqueous buffers (such as saline) is a significant challenge to the clinical translation of injectable cell therapies.^14, 55–57^ In order to assess whether SHIELD encapsulation can safeguard hiPSC-DCNs from cell membrane damage during transplantation, we replicated our standard *in vivo* injection parameters *in vitro* with a spinal cannula (33 gauge needle) and a flow rate of 500 nL/min. At 30 minutes post injection, hiPSC-DCNs delivered in SHIELD experienced significantly less membrane damage compared to hiPSC-DCNs delivered in saline and were statistically similar to hiPSC-DCNs that were not exposed to the injection procedure (Fig. 2E). Having optimized parameters to best promote hiPSC-DCN survival following encapsulation and injection *in vitro*, we next sought to assess the performance of the selected SHIELD formulation *in vivo*.

### SHIELD is cytocompatible and biodegradable within the injured adult cervical spine

Previously, we injected human adipose-derived stem cells encapsulated within our softest SHIELD formulation subcutaneously in uninjured, nude mice and were able to detect fluorescently-labeled material for up to 21 days post injection, but viable cells were no longer present by 3 days post injection.^36^ In a separate study, we injected Schwann cells encapsulated within a stiffer SHIELD formulation (1.25 wt% PNIPAM) into the injured adult cervical spine of Fischer rats. Viable cells were present at the injury site at the end of the study, 28 days post transplantation.^14^

In this study, we chose to test SHIELD (with the C7-RGDS and 0 wt% PNIPAM formulation identified above) in immunodeficient RNU^-/-^ athymic rats for their ability to tolerate human cell xenografts without the need for additional immunosuppression. We first evaluated how long SHIELD persists following injection in this injury model using fluorescently-tagged polymers. Testing SHIELD in this specific injury model is necessary as unwanted immunological and inflammatory tissue responses to biomaterials may be possible given that RNUs are only partially immunodeficient and tissue responses to biomaterials can vary based solely on the site of implantation.^58–60^

Female athymic rats underwent a mid-cervical unilateral mild contusion and received intraspinal injections of fluorescently-labeled C7 alone (*i.e.* with the RGDS ligand but lacking PEG-P1 and hence unable to form a crosslinked hydrogel, 4 μL) or SHIELD (4 μL) two weeks after injury to mimic a subacute therapeutic window (Fig. 3A).^35^ Fluorescence (and therefore material retention) was tracked weekly with an *In Vivo* Imaging System (IVIS) until signal was no longer detectable. When injected into the injured cervical spine, C7 alone was detectable up to 2 weeks post transplantation, whereas SHIELD was detectable up to 5 weeks post transplantation, demonstrating that formation of a crosslinked hydrogel results in slower biodegradation (Fig. 3B-C, **Fig. S4**). No adverse reactions or overt inflammatory responses were evident in animals injected with C7 or SHIELD when compared to an injury-only control. These data suggest that: (1) crosslinking between C7 and PEG-P1 to form SHIELD is necessary for prolonged material retention, and (2) this SHIELD gel formulation can be maintained within the spinal cord for chronic time periods. As immature cortical neurons take time to become functionally active both in development and *in vitro*, we next sought to assess whether SHIELD encapsulation could improve metrics of hiPSC-DCN transplantation and integration after 6 weeks *in vivo*.^61–63^

**Figure 3.**
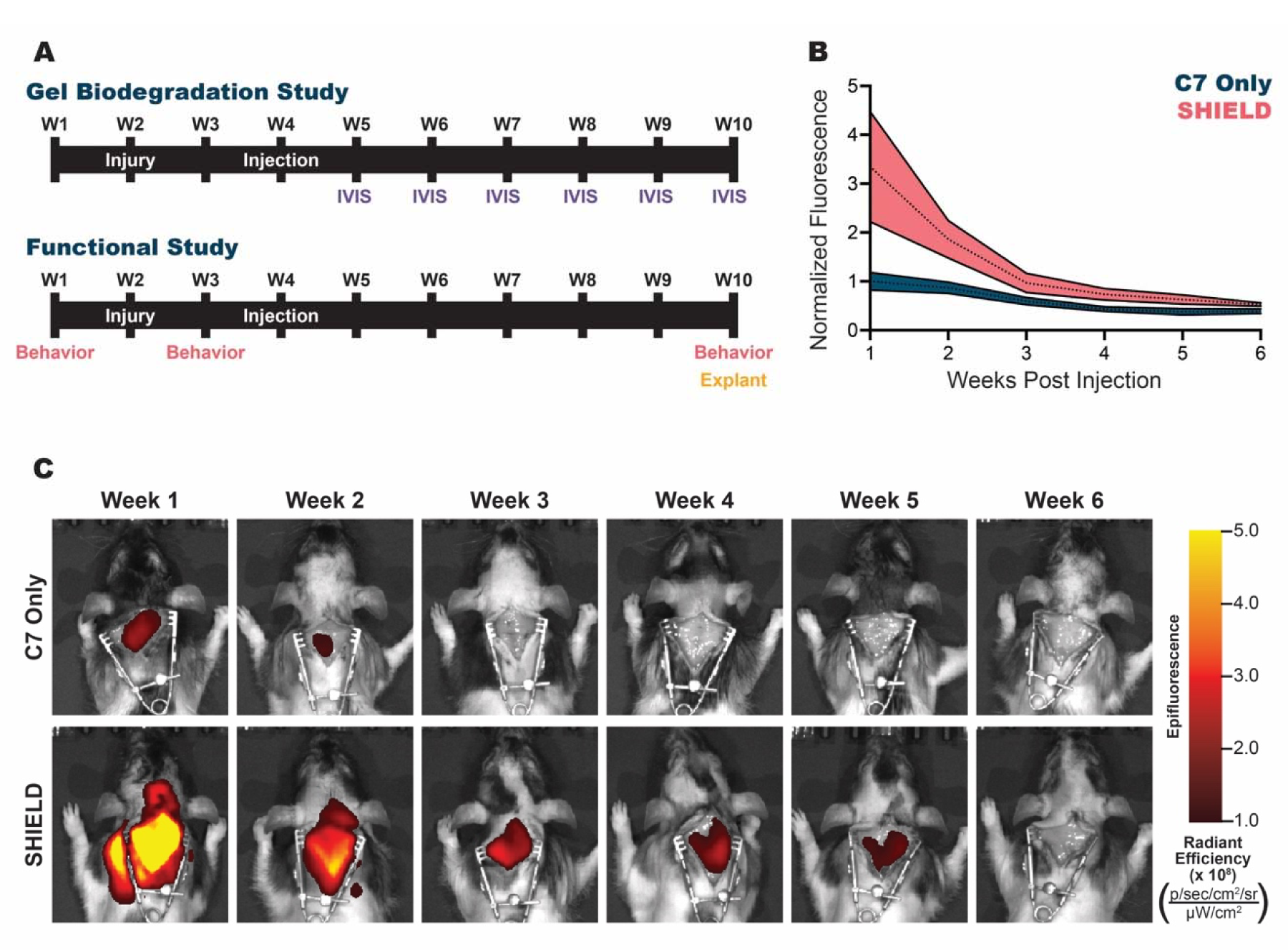
*Crosslinking SHIELD is required for long-term retention within the injured cervical spine.* **A.** Timeline for biodegradation and functional studies depicting surgical procedures (injury, injection; white), live imaging (IVIS; purple), behavioral functional assays (behavior; pink), and endpoints for tissue processing (explant; yellow). **B.** The fluorescence of intraspinally injected cyanine-7 labeled C7 and SHIELD was quantified up to 6 weeks post injection. Data are normalized to C7 at 1 week post injection. Data are mean ± SEM, *n* = 3-4 per condition. **C.** Representative IVIS images from two individual rats intraspinally injected with labeled C7 or SHIELD from 1-6 weeks post injection. In each image, skin was retracted to expose the underlying muscle to avoid masking of signal due to the pigmentation patterns in hooded rats.

### Encapsulating hiPSC-DCNs in SHIELD improves transplantation efficacy

Female homozygous athymic rats received a mid-cervical (C5 level) unilateral mild contusion, were transplanted 2 weeks later, and euthanized 6 weeks post transplantation for graft analysis (Fig. 3A). hiPSC-DCNs delivered in saline dispersed into a larger volume of tissue compared to delivery of hiPSC-DCNs in SHIELD (Fig. 4A). In comparison, hiPSC-DCNs delivered in SHIELD occupied a smaller tissue volume, suggesting that injected cells were successfully retained within the SHIELD gel (Fig. 4A). Despite differences in average tissue volume containing transplanted cells (P = 0.469, Fig. 4A), the total average number of transplanted SC101^+^ cells (human nuclear marker) detected after 6 weeks was statistically similar, regardless of delivery vehicle (P = 0.254, Fig. 4B). For both metrics of graft volume and number of transplanted human cells, the statistical variability was markedly increased for cells delivered in saline compared to SHIELD (F = 16.170, Fig. 4A; F = 4.409, Fig. 4B). In particular, a significant number of rats with saline-based cell delivery had low or no evidence of graft survival, indicative of failed cell delivery due to a combination of reflux and cell death (Figs. 4A, 4B, **S5A**). Taken together, these data demonstrate that delivery in SHIELD resulted in more reproducible transplantation of viable hiPSC-DCNs.

**Figure 4.**
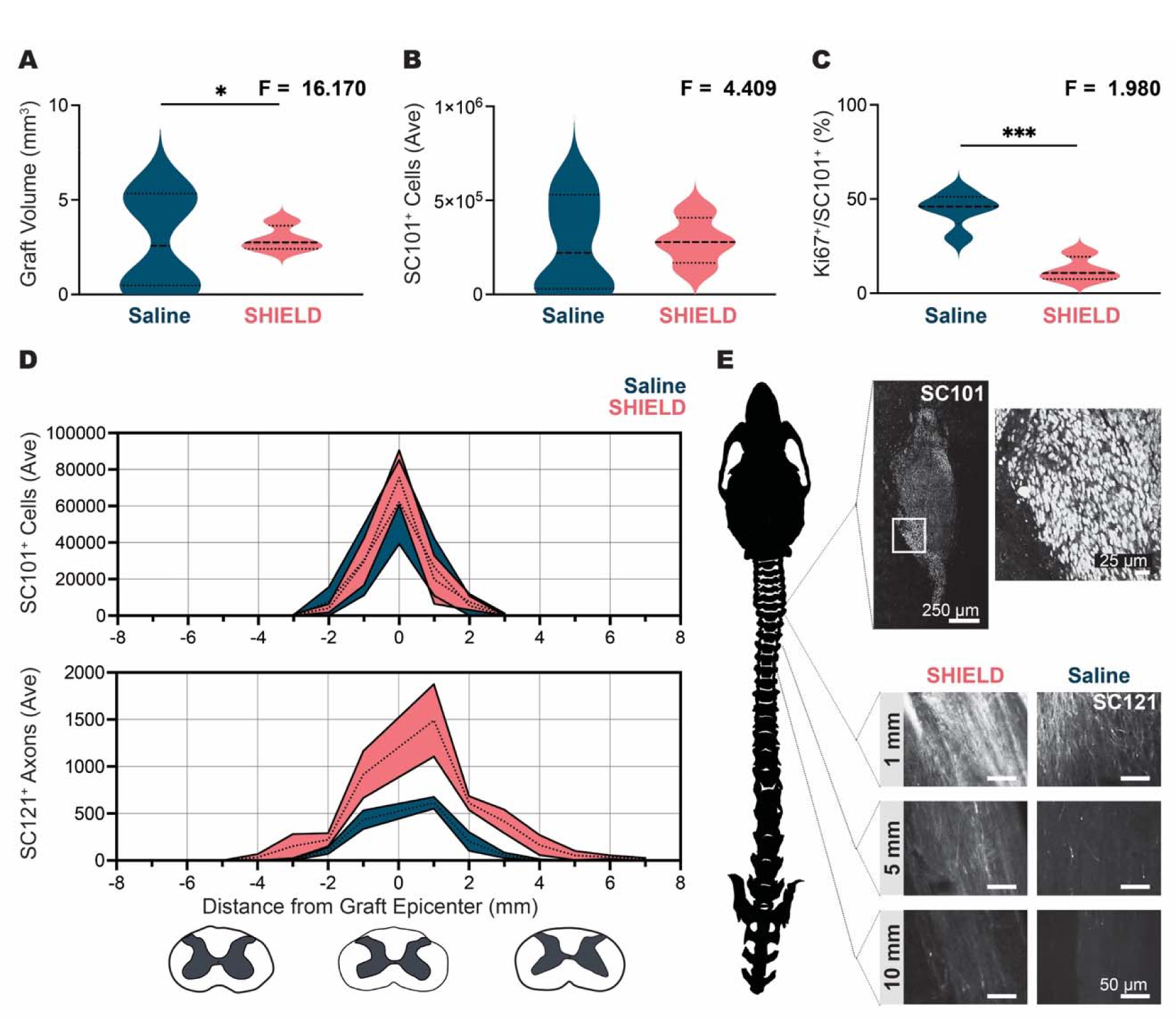
*Encapsulation in SHIELD improves hiPSC-DCN transplantation metrics.* **A.** At 6 weeks post transplantation, graft volume was calculated by measuring the area occupied by SC101^+^ (human nuclear marker) hiPSC-DCNs in serially labeled spinal cord tissue sections. Delivery in saline resulted in a significantly higher graft volume average and larger variability in graft volume when compared to delivery in SHIELD. Data are violin plots; median = dashed line, quartiles = dotted lines; \**P* = 0.0469, F = 16.17; Welch’s t test; *n* = 4. **B.** Quantification of SC101^+^ transplanted cells demonstrated no significant difference in the average number of human cells present within the graft regardless of delivery vehicle. Rats transplanted with hiPSC-DCNs delivered in saline had higher variability in quantified SC101^+^ transplanted human cells. Data are violin plots; median = dashed line, quartiles = dotted lines; *P* = 0.254, F = 4.409; Welch’s t test; *n* = 4. **C.** Quantification of co-labeled Ki67^+^/SC101^+^ human transplanted cells demonstrates a significantly higher number of proliferating human cells in grafts delivered in saline versus SHIELD. Data are violin plots; median = dashed line, quartiles = dotted lines; \*\*\**P* = 0.0005, F = 1.980; Welch’s t test; *n* = 4. **D.** At 6 weeks post hiPSC-DCN transplantation, SC101^+^ human cell bodies (top) and SC121^+^ human projections (bottom) were quantified in 1-mm increments in serial-labeled sections of spinal cord tissue to determine distribution along the continuum of the spinal cord. Distribution and quantity of SC101^+^ human cells were similar regardless of delivery vehicle. Delivery in SHIELD, however, resulted in a higher number of SC121^+^ projections and extension over a longer distance, as appropriate for this cellular phenotype. Data are mean ± SEM. Bottom schematic shows typical spinal cord cross-section at the location of the graft epicenter (C5) and 5 mm rostral (negative) and caudal (positive). **E.** Left: Schematic of the rat cranium and spinal cord vertebrae, depicting location of histological slices. Right top: Representative images of SC101^+^ human cells taken from the transverse plane of the graft epicenter in a rat transplanted with hiPSC-DCNs delivered in SHIELD. Area of inset shown with white box. Right bottom: Representative images of SC121^+^ human projections taken from the transverse plane at locations 1, 5, and 10 mm caudal to the graft epicenter in rats transplanted with hiPSC-DCNs delivered in SHIELD or saline.

Given the known vulnerability of post-mitotic neurons to shear stress, we speculated that delivery in saline may lead to preferential survival of undifferentiated progenitors present in the transplant. Progenitors are proliferative and highly migratory, both of which may contribute to the larger volume of distribution for cells delivered in saline. To evaluate whether proliferation may account for the higher graph volume in a subset of rats transplanted with hiPSC-DCNs delivered in saline, we quantified the percentage of proliferating human cells within the graft (Ki67^+^/SC101^+^) (Fig. 4C, **Fig. S5B**). The fraction of proliferative human cells was higher for transplants delivered in saline compared to those delivered in SHIELD, consistent with preferential survival of undifferentiated progenitors in saline and loss of the much larger population of post-mitotic neurons.

To evaluate whether dispersal or migration may contribute to the higher volume of graft distribution, we quantified SC101^+^ cells in 1-mm increments along the rostral/caudal axis from the graft epicenter (Fig. 4D, top). The border of the cell transplant region typically had clearly defined borders (Fig. 4E, top). We observed that SC101^+^ human cell distribution along the cord was similar regardless of delivery vehicle, with deviation higher when delivered in saline (Fig. 4D, top).

hiPSC-DCNs are pre-patterned *in vitro* to acquire a cortical projection neuron phenotype. One of the hallmarks of this phenotype is long distance, unidirectional axonal growth.^21^ We therefore sought to determine whether transplanted hiPSC-DCNs maintain their subtype identity. To determine if delivery vehicle affected neurite extension, we quantified SC121^+^ (human cytoplasm) projections in 1-mm increments along the rostral/caudal axis from the graft epicenter (Fig. 4D bottom, Fig. 4E, bottom). Delivery in SHIELD resulted in a three-fold higher count of SC121^+^ projections extended from hiPSC-DCNs at the graft epicenter, and these SC121^+^ projections extended for a longer unidirectional distance, as appropriate for this cellular phenotype. These SC121^+^ processes were co-labeled with beta-III tubulin when delivered in SHIELD, indicating commitment to a neuronal phenotype (**Fig. S5C**, **Fig. S6**).

Collectively, these data suggest that SHIELD improves the survival of post-mitotic neurons in the differentiated cell population, consequently promoting more extensive neuritic elongation and integration of the graft into the host spinal cord.

### Improving hiPSC-DCN transplantation engraftment improves functional outcomes

Behavioral outcomes are the most important factor in evaluating the extent of injury and treatment efficacy.^64^ Rats are capable of producing a diverse range of dynamic movements with their forepaws and forelimbs that are comparable to those executed by human and nonhuman primates.^65^ The motor neurons that innervate the forelimb of rats are arranged in columns that run longitudinally and cover multiple segments of the spinal cord.^66^ Cervical SCI eliminates descending inputs to motor neuron pools supplying the muscles of the forelimb ipsilateral to the lesion.^67^ Therefore, we chose three different sensorimotor behavioral tests that collectively assess forelimb function by targeting the different muscle groups of the forelimb (Fig. 5A) to quantitatively compare functional outcomes across our four different treatment groups (Fig. 5B).

**Figure 5.**
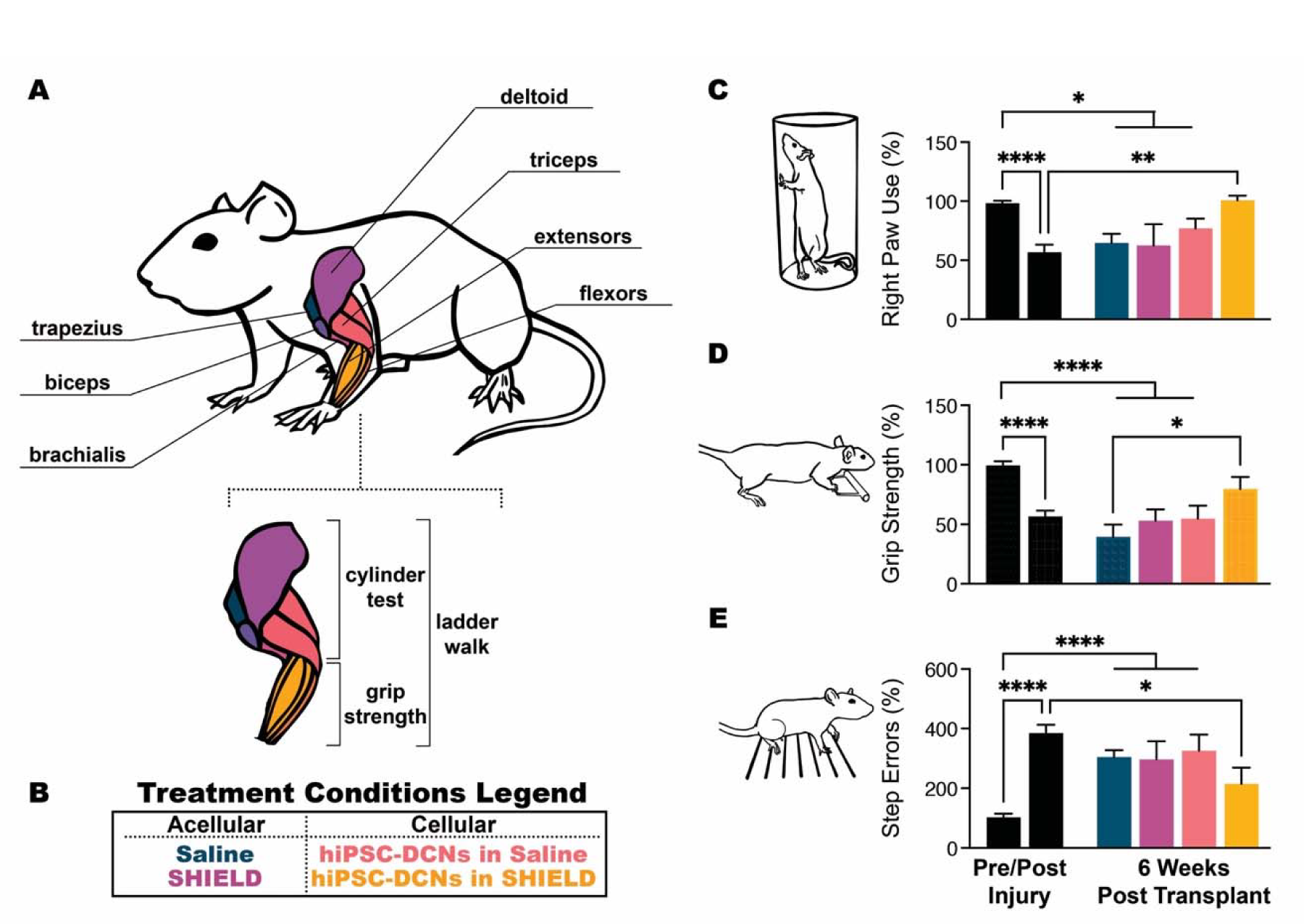
*Transplantation of hiPSC-DCNs encapsulated in SHIELD promotes functional recovery.* **A.** Schematic depicting the major muscles of the forelimb and the behavioral assays that target their function. **B.** Legend depicting the four different treatment conditions for data in panels C-E. **C-E.** All data are mean ± SEM, with one-way ANOVA with Tukey’s multiple comparisons post hoc test. **C.** The cylinder assay quantifies the percentage of right forelimb touches per total touches as a metric of forelimb preference, normalized to pre-injury levels. \**P* = 0.0243, \*\**P* = 0.0027, \*\*\*\**P* < 0.0001. **D.** Grip strength of both forelimbs was quantified using a metered bar, normalized to pre-injury levels. \**P* = 0.0336, \*\*\*\**P* < 0.0001. **E.** Forelimb coordination was assessed with the horizontal ladder walk test. A decrease in coordination presents as an increased percentage of step errors per total steps, normalized to pre-injury levels. \**P* = 0.0217, \*\*\*\**P* < 0.0001.

In behavioral assays measuring forelimb preference (cylinder assay; Fig. 5C), grip strength (Fig. 5D), and skilled locomotion (horizontal ladder walk; Fig. 5E), all rats displayed a significant and permanent decrease in appropriate forelimb sensorimotor function post-injury compared to pre-injury function. By 6 weeks post transplantation, rats that had been transplanted with hiPSC-DCNs delivered in SHIELD displayed improvement in functionality across all three metrics when compared to post-injury performance. In contrast, injured rats that were transplanted with hiPSC-DCNS delivered in saline maintained their post-injury deficit over the course of the study. Control groups of acellular injection with either saline or SHIELD also showed no statistical improvement in function across all three metrics. Collectively, these data suggest that improving hiPSC-DCN engraftment using SHIELD translates to functional improvements.

## Conclusion

Here we presented the design of a delivery vehicle that is customized for transplantation of human cortical projection neurons into the injured cervical spinal cord. We synthesized a family of SHIELD gels with different cell-adhesive ligands and spanning a range of stiffness through the addition of a thermoresponsive polymer. We directed the differentiation of hiPSCs towards a cortical projection phenotype that mimicked hallmarks of human corticogenesis. We quantified hiPSC-DCN survival in the presence of different cell-adhesive ligands and neurite outgrowth in gels of different stiffness and identified the SHIELD variant with the RGDS binding ligand and the most compliant stiffness as being most appropriate for hiPSC-DCN culture *in vitro*. The predominant population of hiPSCs-DCNs prepared for transplant are postmitotic projection neurons with a minor component of undifferentiated progenitors. Neurons encapsulated in SHIELD demonstrated higher viability when compared to cells delivered in saline using an *in vitro* injection assay that was similar to the preclinical transplantation protocol. In a rat model of mid-cervical (C5 level) unilateral mild contusion, SHIELD remained detectable up to 5 weeks post transplantation. When delivered in SHIELD, transplanted hiPSC-DCNs demonstrated significantly improved transplant consistency, with denser transplant regions (*i.e.* a similar number of cells in a smaller tissue volume) that extended a larger number of neuritic projections. Projections from SHIELD-delivered hiPSC-DCNs were also significantly longer, extending into the surrounding spinal cord tissue more than 4 mm rostral and 7 mm caudal to the graft epicenter. Across three functional assays, animals treated with hiPSC-DCNs delivered in SHIELD exhibited significant improvement in behavioral function, while cell transplantation in saline or injection of SHIELD alone resulted in no functional improvement.

Collectively these data demonstrate the potential of customized engineered biomaterials to significantly improve the functional outcomes of cell-based therapy in a preclinical model of cervical spinal cord injury.

## Author Contributions and Acknowledgements

We thank Isabella Ramirez Velez and Eduardo Barrios for their assistance in performing immunofluorescence staining and analysis. We thank Ruby Kemunto Onsongo, Holland Eve Stacey, Sofia Ceva, Ricky Javier Rios, and Syed Ashal Ali for their contributions to animal care.

### Funding

This work was supported by California Institute for Regenerative Medicine RT3-07948 and DISC2-13020 (SCH), National Institutes of Health R01-EB027666 and R01-EB027171 (SCH), Stanford Bio-X IIP (SCH, GWP), Wings for Life WFL-US-020/14 and WFL-US-15/17 (GWP), International Spinal Research Trust STR119 (GWP), Dennis Chan Foundation (GWP), Klein Family Fund (GWP), U.S. Army Medical Research and Materiel Command, Congressionally Directed Medical Research Program W81XWH-18-1-0260 (GWP).

### Author Contributions

V.M.D., L.M.M., G.W.P., and S.C.H. conceived the project. V.M.D., L.M.M., G.W.P., and S.C.H. designed the research. T.D.P., G.W.P., and S.C.H. supervised the research. All authors designed, performed, and/or analyzed all experiments. L.M.M., N.J.B., and M.E.H. synthesized polymer materials. V.M.D., L.M.M., R.A.S., and K.D. performed all surgical procedures. V.M.D., M.E.H., N.J.B., and R.A.S. performed all IVIS imaging. V.M.D., L.M.M., and A.T.W. performed all immunofluorescence staining and imaging of histological samples. V.M.D., L.M.M., and J.M.W. performed cell culture experiments. All authors contributed to manuscript writing.

### Data and materials availability

All data needed to evaluate the conclusions in the paper are present in the paper and/or the Supplementary Materials. Additional data related to this paper may be requested from the authors.

## Materials and Methods

### SHIELD Synthesis and Characterization

The family of SHIELD hydrogels used in these experiments was synthesized and characterized as we have previously described.^14, 36, 37^ SHIELD is comprised of 2 components: (1) C7, a recombinant engineered protein, and (2) a synthetic 8-arm PEG polymer decorated with proline-rich peptides and PNIPAM, a thermoresponsive polymer.

The recombinant C7 proteins contain 7 repeats of the CC43 WW protein binding domain with six repeats of selected fibronectin or laminin-based cell-adhesive domains (RGDS, YIGSR, and IKVAV, and RDGS). C7 variants were expressed in BL21(DE3)pLysS *E. coli* (Invitrogen C606010) as previously described by our group.^14, 38^ Briefly, C7 was cloned into pET-15b plasmids under the control of the T7 promoter, cultured to an optical density at 600 nm of 0.8, and expression was induced via the addition of 1 mM isopropyl β-d-1-thiogalactopyranoside. After 24 hours, the bacteria was centrifuged, harvested, and lysed via multiple freeze-thaw cycles in a lysis buffer (10 mM tris, 1 mM EDTA, 100 mM NaCl). Following the addition of deoxyribonuclease I and 1 mM phenylmethanesulfonyl fluoride protease inhibitor, C7 was purified via affinity chromatography, dialyzed against phosphate buffered saline (PBS), and concentrated by diafiltration. C7 protein purity was confirmed via SDS-polyacrylamide gel electrophoresis, Western blotting, and amino acid sequencing. PEG-P1 and PEG-P1-PNIPAM were synthesized as we have previously described.^14, 36, 68^ Briefly, PNIPAM endcapped with a thiol group was synthesized using reversible addition–fragmentation chain transfer polymerization and conjugated to the 8-arm PEG vinyl sulfone (Nanocs PG8A-VS-20k) via a Michael-type addition reaction. The PNIPAM conjugation reaction was altered to modify either 0.5 or 1 arm of the PEG-vinyl sulfone while the unreacted 7 arms were further reacted with excess P peptide (EYPPYPPPPYPSGC, 1563 g/mol; GenScript Corp.). Conjugation reactions were confirmed via ^1^H nuclear magnetic resonance. The PEG-P-PNIPAM copolymer solution was lyophilized, washed with chloroform, and dialyzed against deionized water (pH 7.4) to remove unreacted PNIPAM and P.

SHIELD gels were prepared by adding PBS to reach a 10% (w/v) C7 and a 10% (w/v) PEG-P or PEG-P-PNIPAM solution. For gel fabrication, each WW domain in C7 was treated as one C unit, and each pendant P peptide group in the PEG-P-PNIPAM copolymer was treated as one P unit and components were mixed to achieve a C:P ratio of 1:1 and final polymer concentration of 10% (w/v). To fabricate the three different SHIELD mechanical formulations (0%, 1.25%, 2.5% PNIPAM), appropriate amounts of 10% (w/v) PEG-P and/or PEG-P-PNIPAM were added to the 10% (w/v) C7 solution. The final concentration of cell-adhesive peptides is estimated to be 6.5 mM.^14^ Dynamic oscillatory rheology experiments were performed on a stress-controlled rheometer (AR-G2, TA Instruments) using a 20-mm diameter cone plate geometry. Samples were loaded and a humidity chamber was utilized to prevent dehydration. Frequency sweeps from 0.1 to 10 Hz at 25° and 37°C were performed at 1% constant strain to obtain storage moduli (*G*′) and loss moduli (*G*″).

### hiPSC Culture

For these experiments, the human iPSC line HuF5.3 was used.^35, 69^ hiPSCs were plated in 6-well plates (Corning 3516) that had been coated with Matrigel hESC Qualified Matrix (Corning 354277) using the thin-coating protocol, per the manufacturer’s instructions. Protein concentration of Matrigel was kept consistent throughout the experiments. Undifferentiated hiPSCs were maintained in an E8 media cocktail (Gibco A1517001) with daily media changes. Colony morphology was assessed daily for cobblestone appearance, uniform and tightly packed cells, and clearly defined borders. Colonies that did not meet these criteria were manually discarded using a sterile P200 pipette tip under microscopic observation prior to passaging. hiPSCs were not maintained on an antibiotic/antimycotic cocktail but spent media was regularly tested for mycoplasma (Lonza LT07-318).

hiPSCs were passaged 2 to 3 times weekly. Remnant media was aspirated and the culture was gently rinsed twice with 1X DPBS without calcium or magnesium (Gibco 14190144). EDTA (0.5 mM in DPBS; Corning 46034CI) was added as a chelating agent for 3-5 minutes at 37°C. Once changes in colony morphology was observed (edges of colonies curling up, lace-like appearance), the EDTA was aspirated and E8 media was used to dislodge adherent colonies from the plate. The collected colonies were gently triturated (care taken to avoid a single cell suspension) and spun at 200 x g for 3 minutes at room temperature. The cell pellet was then gently resuspended in E8 media supplemented with Y27632, a Rho-associated protein kinase inhibitor (3 μM; Stemgent 04-0012) and plated on Matrigel-coated 6-well plates. In subsequent media changes, Y27632 was not included in the E8 media cocktail.

### Directed Differentiation of hiPSC-DCNs

hiPSC cultures maintained in E8 media were chosen for differentiation if the culture reached 70-80% confluency while maintaining standard, uniform morphology across colonies. At this point, the hiPSC culture was differentiated towards a deep cortical neuron-enriched phenotype using an adaptation of the dual SMAD inhibition protocol.^35, 39, 70^ E8 media was aspirated and replaced with a Differention Media cocktail designed to stimulate cortical neurogenesis via the removal of FGF2 consisting of: DMEM/F12 (48.25%; Gibco 11330-032), Neurobasal-A (48.25%; Gibco 10888022), B-27 (1%; Gibco 17504044), GlutaMAX (1%; Gibco 35050061), MEM non-essential amino acids (1%; Gibco 11140050), and N-2 (0.5%; Gibco 17502048). To drive the cultures towards a neuroepithelium fate, small molecules were added to the Differentiation Media to block TGFβ and BMP signaling. LDN-193189 is a cell-permeable BMP inhibitor of ALK2 and ALK3 while SB431542 inhibits ALK4, ALK5, and ALK7. From Days 0-9 of differentiation, SB435142 (10 μM; Stemgent 04-0010) was added. From Days 0-19 of differentiation, LDN-193189 (500 nM; Stemgent 04-0074) was added. By Day 12, the differentiating cultures are predominantly identified as early dorsal neural progenitors. At this point, the cultures were dissociated (as described above, “*hiPSC Culture*”) and replated onto 6-well plates (Corning 3516) sequentially coated with a heavy molecular weight poly-d-lysine (50 μg/mL diluted in 0.1M Borate Buffer; Sigma P1024) and laminin (10 μg/mL diluted in DPBS; Roche 11243217001). To rostrally shift the cultures towards a dorsal telencephalic fate, Wnt signaling was inhibited (via the prevention of palmitylation of Wnt proteins by Porcn) from Days 12-19 using the small molecule Wnt-C59 (5 nM; Tocris 5148). After 35 days of differentiation, the hiPSC-DCN cultures were papain-dissociated for characterization, *in vitro* injection studies, or *in vivo* transplantation using an adaptation of a well-established neural cell isolation method (Worthington 9035-81-1).^71^ As with our undifferentiated hiPSC cultures, differentiating hiPSC-DCNs were not maintained on an antibiotic/antimycotic cocktail but spent media was regularly tested for mycoplasma (Lonza LT07-318).

### In Vitro Viability Studies

For *in vitro* injection studies, hiPSC-DCNs were papain-dissociated after 35 days of differentiation and resuspended to achieve a final cell density of 75,000 cells/μL. hiPSC-DCNs were encapsulated in an equal volume of (1) a C7 variant (RGDS, YIGSR, IKVAV, or RDGS), (2) a SHIELD variant with differing concentrations of PNIPAM (0.0%, 1.25%, or 2.5%), or Hank’s Balanced Salt Solution and loaded into a 10 μL Hamilton Model 701 RN syringe (Hamilton 7635-01) outfitted with a 30 gauge style 4 needle (Hamilton 7803-07). hiPSC-DCNs were injected onto square 22 mm x 22 mm coverslips using the same injection parameters (500 nL/min with a 4 min rest) as used *in vivo* for transplantation with an automated microinjection syringe pump. After a 30 minute incubation period at 37°C, cell viability/cytotoxicity was assessed via the LIVE/DEAD kit (Invitrogen L3224), following the manufacturer’s instructions. The coverslips were imaged on a Leica Thunder inverted microscope at 10X magnification and analyzed via ImageJ.

### Immunocytochemistry

Encapsulated hiPSC-DCNs were fixed with 4% paraformaldehyde in PBS at 37°C for 30 minutes. Samples were permeabilized with 0.02% Triton X-100 (Sigma 93443) in PBS for 1 hour at room temperature and subsequently blocked with 5% bovine serum albumin (ThermoFisher Scientific 37525) and 10% donkey serum (Lampire 7332100) in Triton X-100 in PBS for 1 hour at room temperature. The samples were then incubated overnight at 4°C with primary antibody (beta-tubulin III 1:500; STEMCELL Technologies 60052) diluted in blocking buffer. The following day, the samples were washed 3 times with PBS and then incubated for 4 hours at room temperature in the dark with secondary antibody (AlexaFluor488 donkey anti mouse 1:400; Jackson 715-545-151) diluted in blocking buffer. Samples were washed thoroughly with PBS and mounted using VECTASHIELD HardSet Antifade Mounting Medium (Vector Laboratories H-1400-10). Samples were imaged using a Leica SPE confocal microscope at 10X magnification and analyzed via ImageJ.

### Animals

Female adult athymic RNU rats (Crl:NIH-Foxn1^rnu^; Charles River Laboratories) were used in this study for their incomplete immunodeficiency, which allows for xenografting. Rats were housed two per cage in our animal facility under a 12/12 hour light/dark cycle and were provided with food and water *ad libitum*. All animal procedures were performed at Stanford University, under an approved Administrative Panels on Laboratory Animal Care Protocol.

### Cervical Spinal Contusion Surgery

A mild unilateral contusion was induced at the 5^th^ cervical level as we have previously described.^14^ At 8 weeks old, animals were weighed and placed under inhalation anesthesia using isoflurane (3%, 1.5% O_2_) before surgery. Once under surgical-level anesthesia (confirmed via lack of withdrawal reflex), rats were shaved around the neck and back region, and aseptically prepared with 70% ethanol and betadine solution. A dorsal, midline skin incision was made, and the skin and underlying muscle layers teased apart from the 3rd cervical (C3) segment to the 2nd thoracic (T2) segment (location determined by counting vertebrae and using the distinctive T2 spinous dorsal process as a landmark). The spinous processes were exposed and a C5 dorsal laminectomy performed to expose the underlying spinal cord. The 4^th^ and 6^th^ cervical vertebrae were cleaned of any tissue and stabilized with clamps to place the exposed C5 spinal cord on a level plane and to minimize movement due to breathing. The animal was then positioned under an impactor device (Infinite Horizons) and a mild 75 kdyne contusion injury was delivered to the right side of the cord. After injury, the individual muscle layers were sutured and the skin closed using 9 mm wound closure clips. Rats were placed in clean, temperature-controlled cages and monitored until walking and righting abilities were regained. At the beginning of surgery and twice daily for 3 days post-operatively, rats received subcutaneous injections of analgesic (buprenorphine HCL, 0.14 mg/kg), antibiotics (penicillin, 115 mU/kg), and saline (to prevent dehydration and normalize blood pressure). Throughout the study, all animals were inspected daily for healing, body condition, and pain with veterinary care given as deemed appropriate.

### Transplantation Surgery

Transplantation occurred 14 days after the SCI surgery as we have previously described.^14^ Injured rats were randomized into one of the following treatment groups: (1) injury only (sham injection - needle insertion only), (2) saline only, (3) SHIELD only, (4) 150,000 hiPSC-DCNs in saline, or (5) 150,000 hiPSC-DCNs in SHIELD. The laminectomy and underlying cervical spine were surgically re-exposed as in the initial SCI surgery and the animal was positioned within a stereotactic set-up to ensure that the exposed spinal cord was level. Treatments were injected into the contusion site (visualized by a dark cavity) after being loaded into a 10 μL Hamilton Model 701 RN syringe (Hamilton 7635-01) outfitted with a 30 gauge style 4 needle (Hamilton 7803-07). Injections were performed using a microinjection apparatus (Quintessential Stereotaxic Injector, Stoelting Co.). The total volume of each injection was 4 μL delivered at a rate of 500 nl/min with a 4 minute additional rest time (syringes left in place) to minimize reflux following needle withdrawal. The individual muscle layers and skin were sutured and post-operative care provided (as described above, “*Cervical Spinal Contusion Surgery*”).

### IVIS Spectrum Imaging

C7 was conjugated using our previously described protocols.^36^ First, Cyanine7 (Cy7) NHS Ester (Lumiprobe 15020) was dissolved in dimethyl sulfoxide (DMSO). C7 protein was diluted to 1 wt% in PBS and pH adjusted to 8.0 using NaOH. The dissolved Cy7 NHS Ester was added to the C7 solution at a 0.1 molar equivalence with the C7 lysine groups and mixed on a rotator for 24 hours. Unreacted Cy7 NHS Ester was filtered out using a spin filter (MWCO 10 kDa, Millipore Amicon Ultra-0.5). Fluorophore functionalized C7 was mixed with unfunctionalized C7 at a 1:1.5 ratio. At 2 weeks post cervical SCI, rats were injected with 4 μL of SHIELD or C7. Each animal was imaged using an In Vivo Imaging System (IVIS Lago) over a series of 42 days. Prior to imaging, rats were anesthetized and a longitudinal incision was surgically induced to retract the skin away from the underlying muscle layers above the cervical spine. As hooded rats have variable dorsal pigmentation patterns and dark pigmentation obstructs visualization of the Cy7 signal, retracting the skin normalizes for differences amongst individual rats. Rats were then positioned within the IVIS imaging apparatus and imaged with an automatic exposure time, excitation wavelength of 740, emission wavelength of 790 (binning; medium, F/stop; 2.0). Following imaging, the skin incision was closed using 9 mm wound closure clips and post-operative care was provided (as described above, “*Cervical Spinal Contusion Surgery*”). Average radiant efficiency was quantified and normalized to day 7 across animals.

### Behavioral Testing

#### Cylinder Test

Forelimb preference was performed as previously described.^64, 72^ Briefly, forelimb preference was measured by placing the rat in a clear Plexiglass cylinder and filming spontaneous exploratory behavior for 3 minutes. Mirrors were placed at an angle behind the cylinder so that the forelimbs could be viewed at all times and positions. The number of times the rat touched its left or right forepaw to the cylinder was tallied and analyzed for changes in symmetry from pre-injury.

#### Grip Strength

Forelimb grip strength was measured on the animals as previously described.^14, 64^ Briefly, rats’ gripping reflex was triggered by allowing the animal to hold on firmly to a metered bar (TSE Systems). The rat was then pulled backwards with a continuous movement, in line with the attachment axis of the grip and the measured value recorded. This was repeated 5 times per animal per timepoint, upon which the lowest and highest values were dropped and the remaining 3 values averaged together. Grip strength was then reported as a change from pre-injury baseline.

#### Horizontal Ladder Walk Test

Skilled walking was measured as previously described.^14, 64, 73^ Briefly, rats were placed within a Plexiglas alleyway with metal rungs irregularly spaced. Animals were recorded (Sony HDR-CX675 camera) walking across the rungs, and the number of correct and incorrect forelimb steps was recorded by blinded observers. The percentage of missed steps was calculated by dividing the number of incorrect steps by the total (correct and incorrect steps) number of steps and multiplying the quotient by 100. Data was then reported as a change from pre-injury baseline.

### Euthanasia and Tissue Processing

At the end of the experimental timeline (6 weeks post transplantation) rats were euthanized and spinal cord tissue was excised for immunohistochemical analysis. Rats received a lethal dose of Beuthanasia® (150 mg/kg IP). Once surgical-level anesthesia was reached, rats were placed on a perfusion stage and positioned to allow for exposure of the peritoneal cavity. Rats were transcardially perfused with ice cold 300 mL of PBS followed by 300 mL of fresh 4% paraformaldehyde (PFA; Sigma P6148) solution. Following transcardial perfusion and fixation, spinal cords were immediately dissected and immersion-fixed in PFA for 24 hours at 4°C. Spinal cords were then cryoprotected in a 30% sucrose (w/v in PBS; Sigma S7903) solution until the tissue sunk to the bottom of the collection tubes. Spinal cords were then embedded in a 10% porcine gelatin (w/v in PBS; Sigma G2500) solution and immersion-fixed in PFA for 24 hours at 4°C. The gelatin-embedded spinal cords were then cryoprotected in a 30% sucrose solution until the blocks sunk to the bottom of the collection tubes. Tissue blocks were then trimmed and sectioned in the transverse plane (60 μm thickness) using a freezing sledge microtome and stored in PBS until use.

### Immunohistochemistry

Tissue sections were blocked for 2 hours at room temperature with 10% normal donkey serum (Lampire 7332100), 0.1% Triton-X (Sigma 93443), and a PBS solution and then incubated with the primary antibody diluted in this blocking buffer while rocking for 48 hours at 4°C. Sections underwent 3 washes with blocking buffer followed by incubation with the secondary antibody diluted in blocking buffer for 4 hours at room temperature in the dark. Sections were washed with a PBS solution 3 times and mounted on microscope slides with ProLong Diamond Antifade Mountant (ThermoFisher P36961) and left to cure for 24 hours at room temperature in the dark. Once properly cured, slides were sealed with nail polish and imaged. The following primary antibodies were used: beta-tubulin III (STEMCELL Technologies 60052; 1:1000), GFAP (Dako GA524; 1:2000), Stem101 (Takara Y40400; 1:100), Stem121 (Takara Y40410; 1:1000), and Stem123 (Takara Y40420; 1:500). The following secondary antibodies were used at a 1:800 dilution: Donkey anti mouse AlexaFluor 488 (Jackson 715-545-151), Donkey anti mouse Cy3 (Jackson 715-165-150), Donkey anti mouse Cy5 (Jackson 715-175-150), Donkey anti rabbit Cy3 (Jackson 711-165-152), and Donkey anti rabbit Cy5 (Jackson 711-175-152).

### Statistics

All data are presented as mean ± SD (*in vitro*) or mean ± SEM (*in vivo*), and statistical analysis was performed using GraphPad Prism software. Statistical comparisons on *in vitro* cell viability and neurite extension were performed by one-way analysis of variance (ANOVA) with Dunnet’s multiple comparisons post hoc test. Statistical comparisons on in *vivo* graft survival metrics were performed by unpaired, two-tailed *t* test with Welch’s correction. Statistical comparisons on functional behavior assays were performed by one-way ANOVA with Tukey’s multiple comparisons post hoc test. In all statistical analyses, a significance criterion of \**P* ≤ 0.05, \*\**P* ≤ 0.01, \*\*\**P* ≤ 0.001, and \*\*\*\**P* ≤ 0.0001 was used.

