## Supplemental Materials for "Custom-engineered hydrogels for delivery of human iPSC-derived neurons into the injured cervical spinal cord"

**List of Supplemental Materials**

**Table S1.** *Amino acid sequences included in the C7 variants and P peptide.*

**Figure S1.** *Heterodimeric binding of C7 and PEG-P1 forms a gel.*

**Figure S2.** *Directed differentiation protocol results in high neuronal purity.*

**Figure S3.** *hiPSC-DCNs differentially respond to biochemical and biomechanical cues.*

**Figure S4.** *Material retention is prolonged for SHIELD compared to uncrosslinked C7 protein.*

**Figure S5.** *hiPSC-DCN engraftment is improved when delivered in SHIELD.*

**Figure S6.** *Transplanted hiPSC-DCNs maintain neural phenotype.*

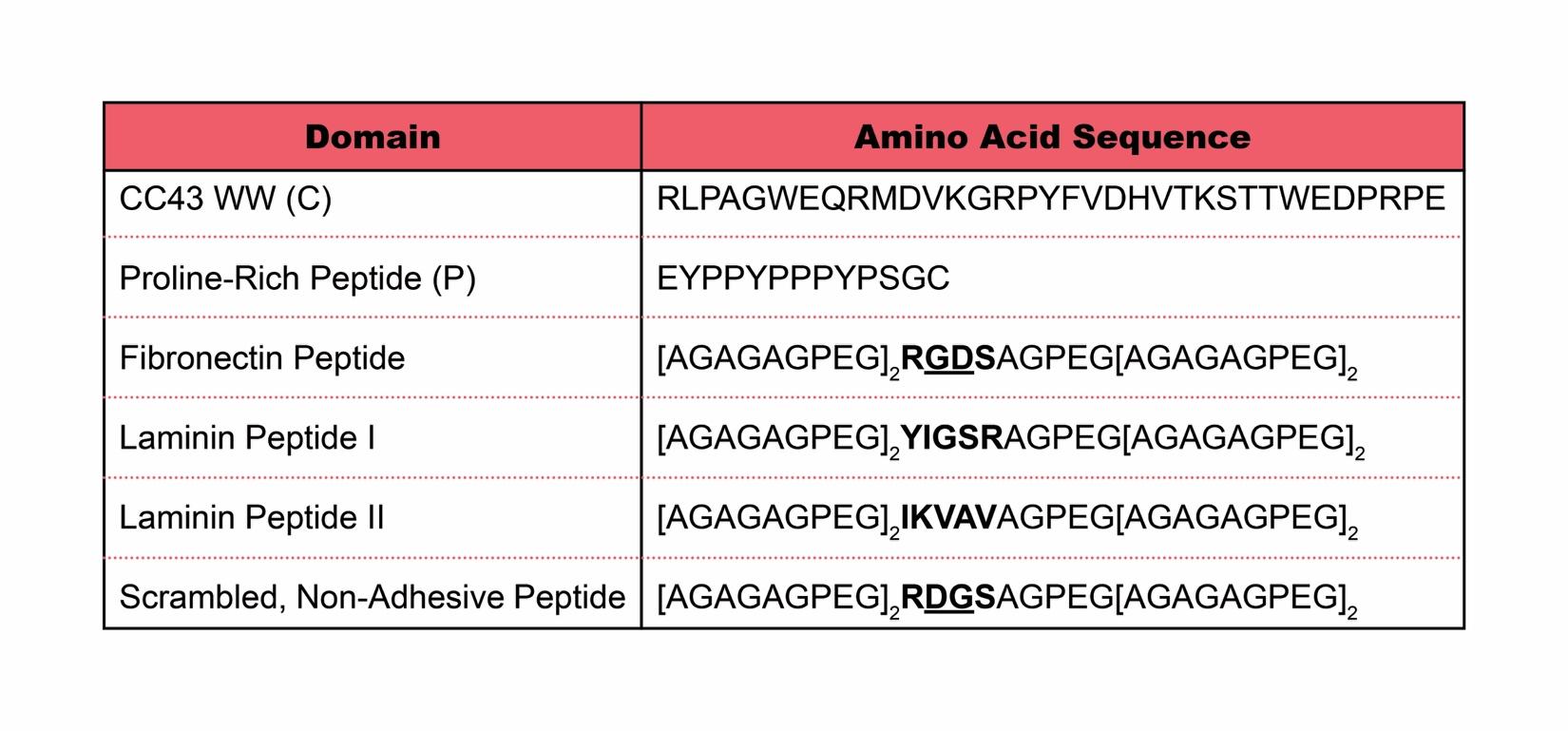

**Table S1.** *Amino acid sequences included in the C7 variants and P peptide.* The C7 variants include alternating repeats of the CC43 WW domain (C) and one of the listed cell-adhesive spacer variants (Fibronectin Peptide, Laminin Peptide I, Laminin Peptide II) or the control spacer sequence (Scrambled, Non-Adhesive Peptide). The multi-arm PEG is conjugated with the proline-rich peptide (P) to form PEG-P1, which can be further modified to include an optional PNIPAM copolymer.

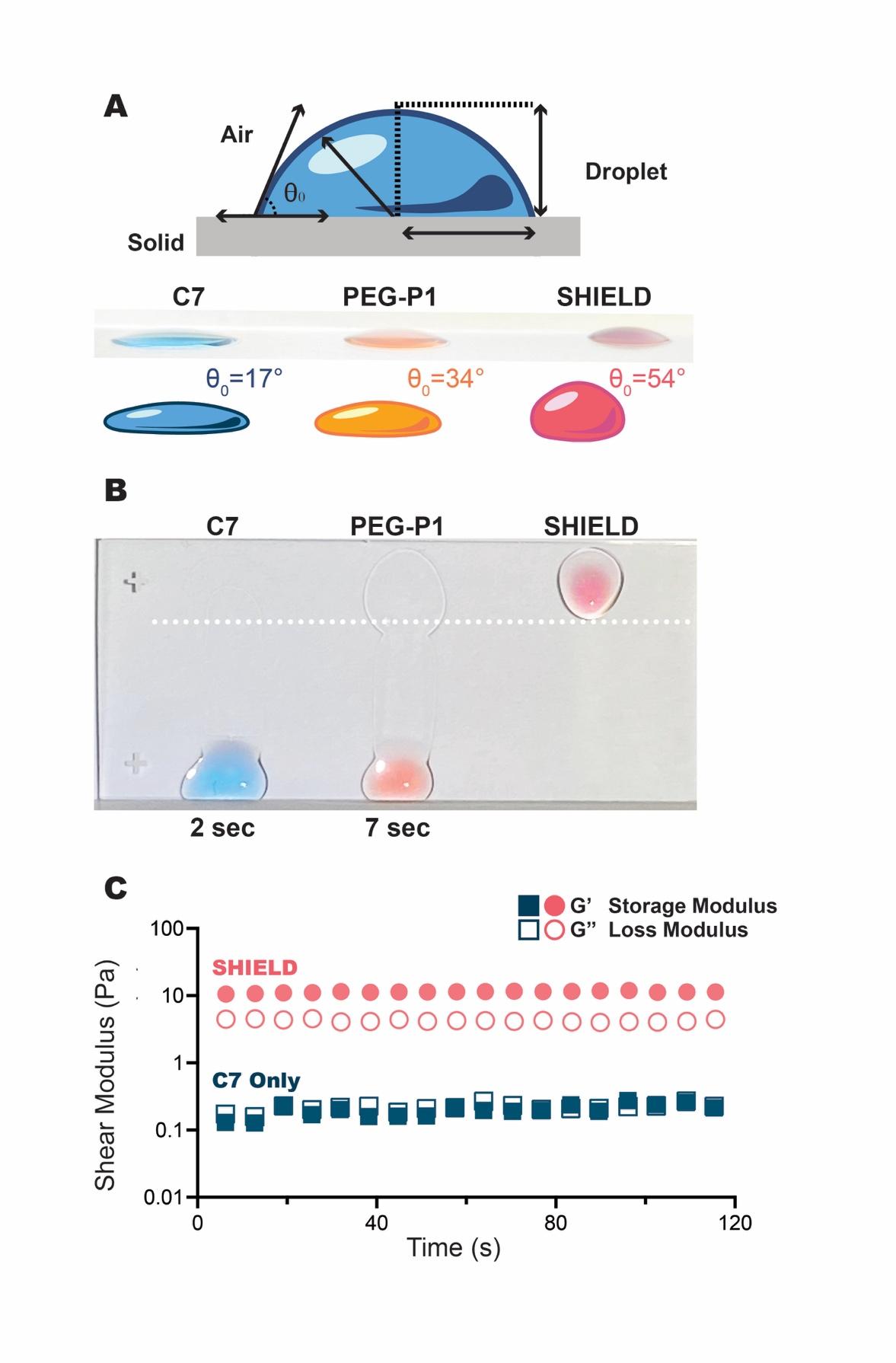

**Figure S1.** *Heterodimeric binding of C7 and PEG-P1 forms a gel.* **A.** The contact angles of C7, PEG-P1 (with 0 wt% PNIPAM), and SHIELD droplets were measured as a metric of wettability. Contact angles lower than 90º are indicative of hydrophilicity. Food coloring added for enhanced visualization. **B.** Ejection of individual (C7, PEG-P1) or pre-mixed components (SHIELD) onto vertical glass slides demonstrating the sol-gel phase transition upon heterodimeric binding of the peptide domains of the individual components at room temperature. Food coloring was added for enhanced visualization. Individual components flow as liquids, covering the length of the slide in seconds; SHIELD solidifies into a weak gel and resists flow. **C.** Representative time sweep for C7 and SHIELD indicating that a gel is formed only when components are mixed to form SHIELD.

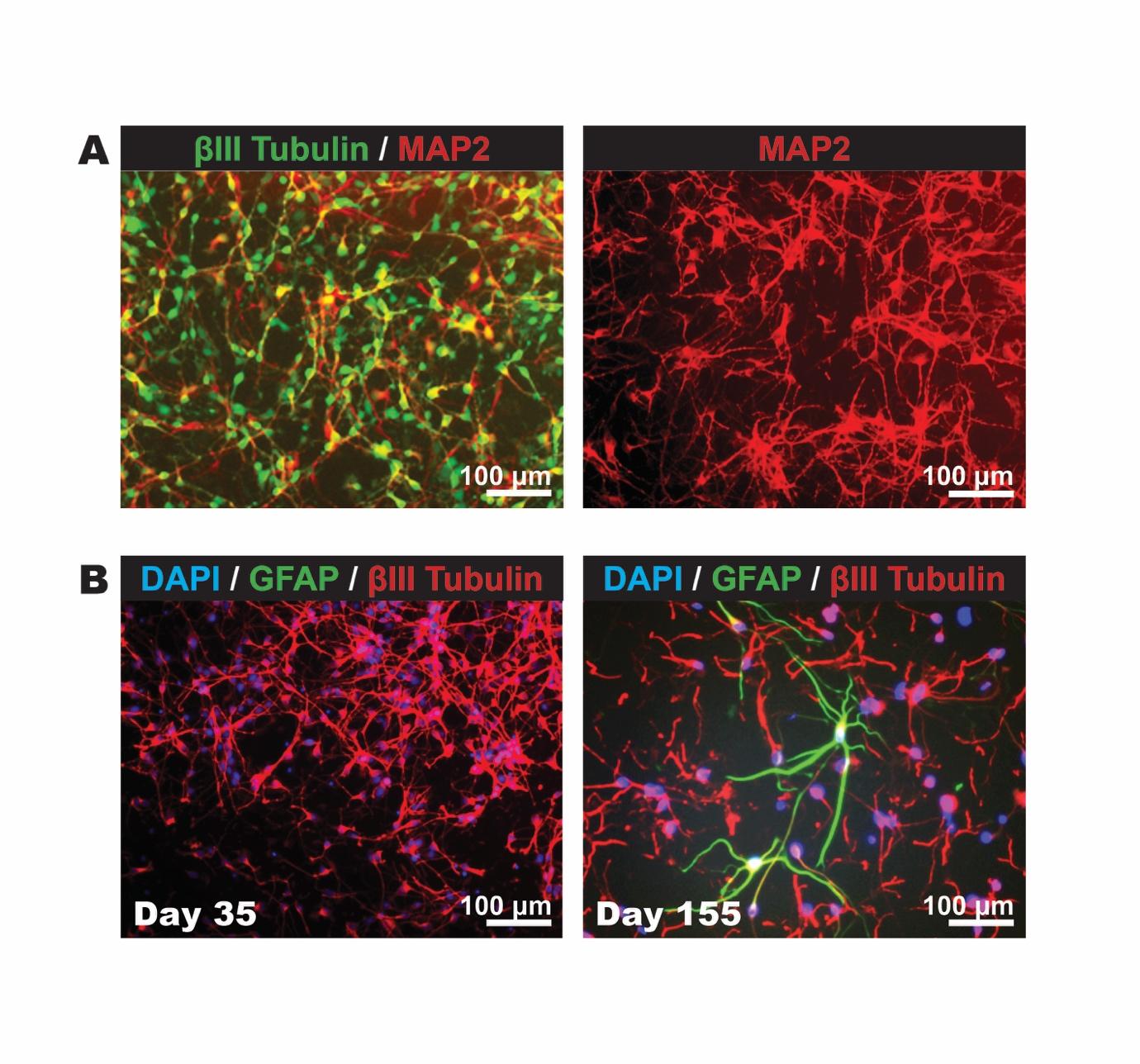

**Figure S2.** *Directed differentiation protocol results in high neuronal purity.* **A.** By 35 days of directed differentiation, most cells within the culture have committed to a neuronal lineage. Extensive neurite networks are observed, and the culture shows robust expression of βIII tubulin (pan-neuronal) and MAP2 (pan-neuronal). **B.** In the developing nervous system, gliogenesis occurs later than neurogenesis. Differentiating hiPSC cultures were imaged after 35 days and after 155 days for nuclei (DAPI), neuronal (βIII tubulin) and astrocytic (GFAP) markers. At 35 days (transplantation stage), the cultures infrequently express GFAP. At longer culture times, some GFAP staining is observed.

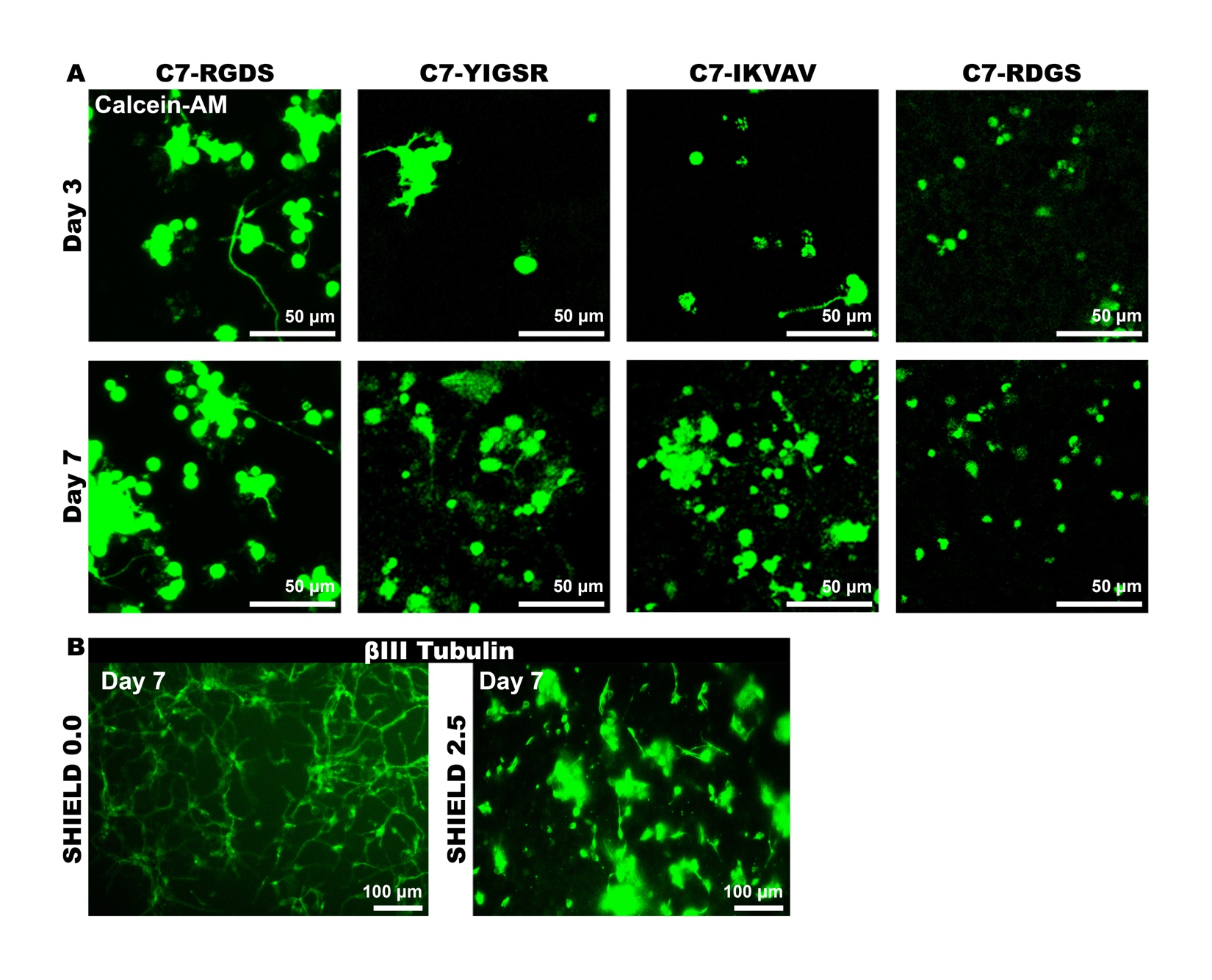

**Figure S3.** *hiPSC-DCNs differentially respond to biochemical and biomechanical cues.* **A.** hiPSC-DCNs were cultured in SHIELD gels formulated with four different C7 variants for 7 days. Calcein-AM^+^ staining demonstrates highest hiPSC-DCN survival and neurite outgrowth in SHIELD with C7-RGDS and lowest with C7-RDGS, a scrambled non-adhesive control. **B.** hiPSC-DCNs were cultured in SHIELD variants (all with C7-RGDS) of increasing stiffness (0.0% and 2.5% w/v PNIPAM) for 7 days. βIII tubulin^+^ staining demonstrates highest hiPSC-DCN outgrowth in the SHIELD formulation with 0% w/v PNIPAM.

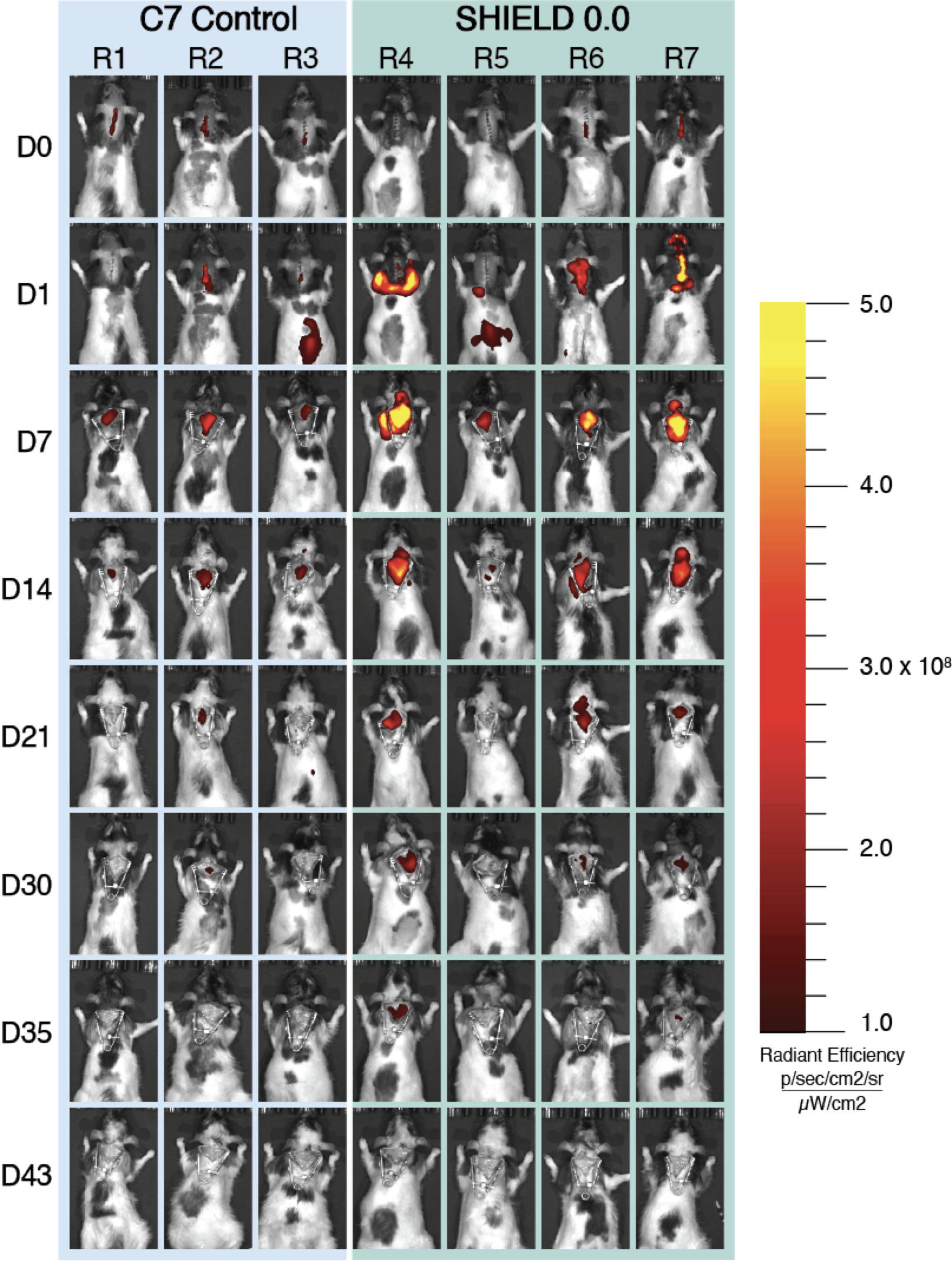

**Figure S4.** *Material retention is prolonged for SHIELD compared to uncrosslinked C7 protein.* IVIS images show the difference in material retention between the uncrosslinked protein C7 alone (n = 3) and the SHIELD 0.0 hydrogel (n = 4) over 43 days in a rat model of cervical spinal cord injury.

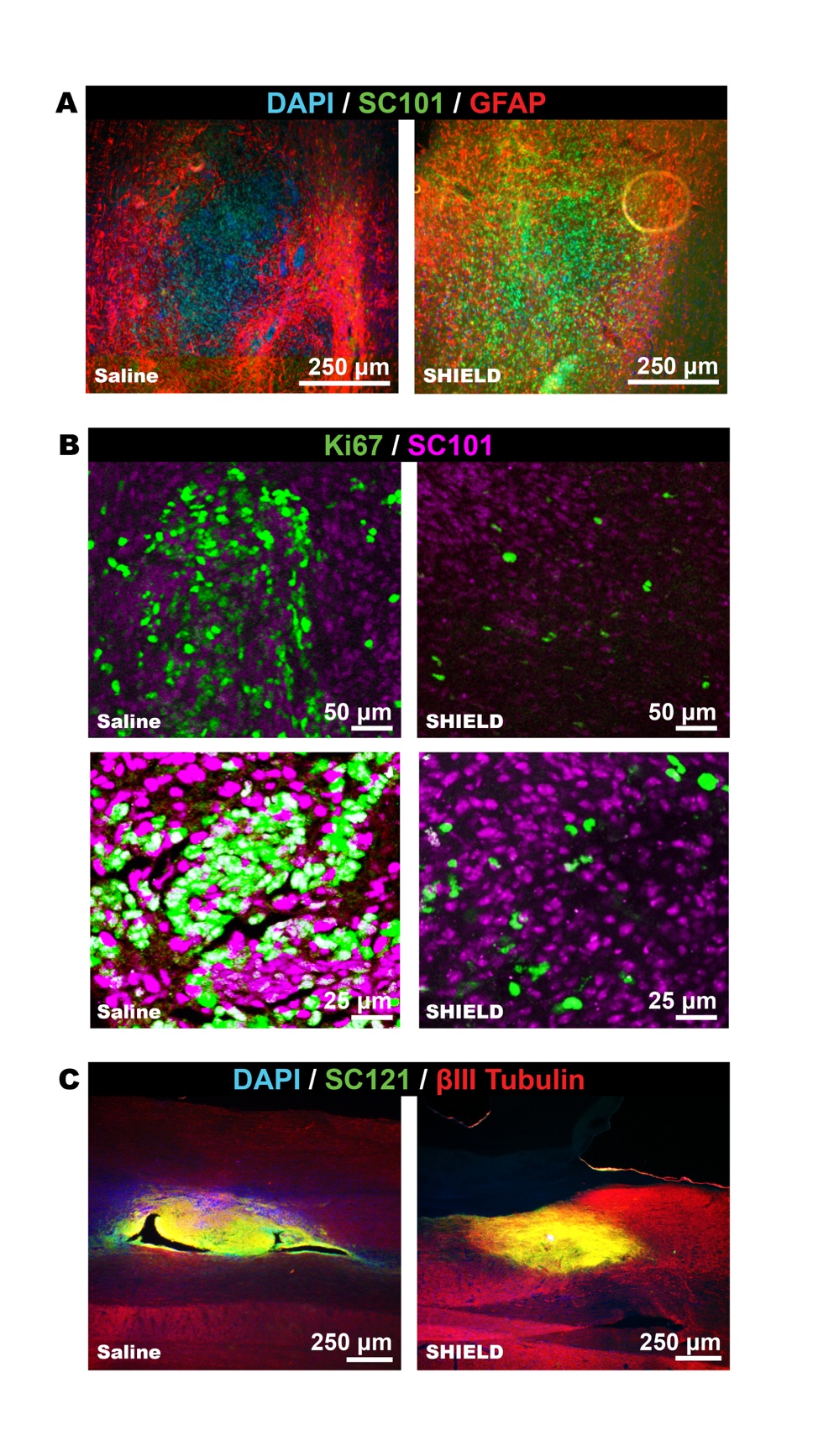

**Figure S5.** *hiPSC-DCN engraftment is improved when delivered in SHIELD.* **A.** Representative images taken from the graft epicenter in rats transplanted with hiPSC-DCNs delivered in either saline or SHIELD. Compared to delivery in saline, SHIELD encapsulation improves hiPSC-DCN engraftment (SC101^+^ human nuclei, green) and reduces astrogliosis (GFAP, red). **B.** Representative images taken from the graft epicenter. When delivered in saline, a higher percentage of transplanted human nuclei (SC101^+^, purple) are also Ki67^+^ (green, white overlay), indicative of proliferation. **C.** Representative images from the graft epicenter. A higher fraction of hiPSC-DCN processes (SC121^+^ human cytoplasm, green) are co-labeled with βIII tubulin^+^ (red) when delivered in SHIELD, indicating commitment to a neuronal phenotype. βIII tubulin^+^ signal is more robust caudal to the graft in rats transplanted with hiPSC-DCNs in SHIELD.

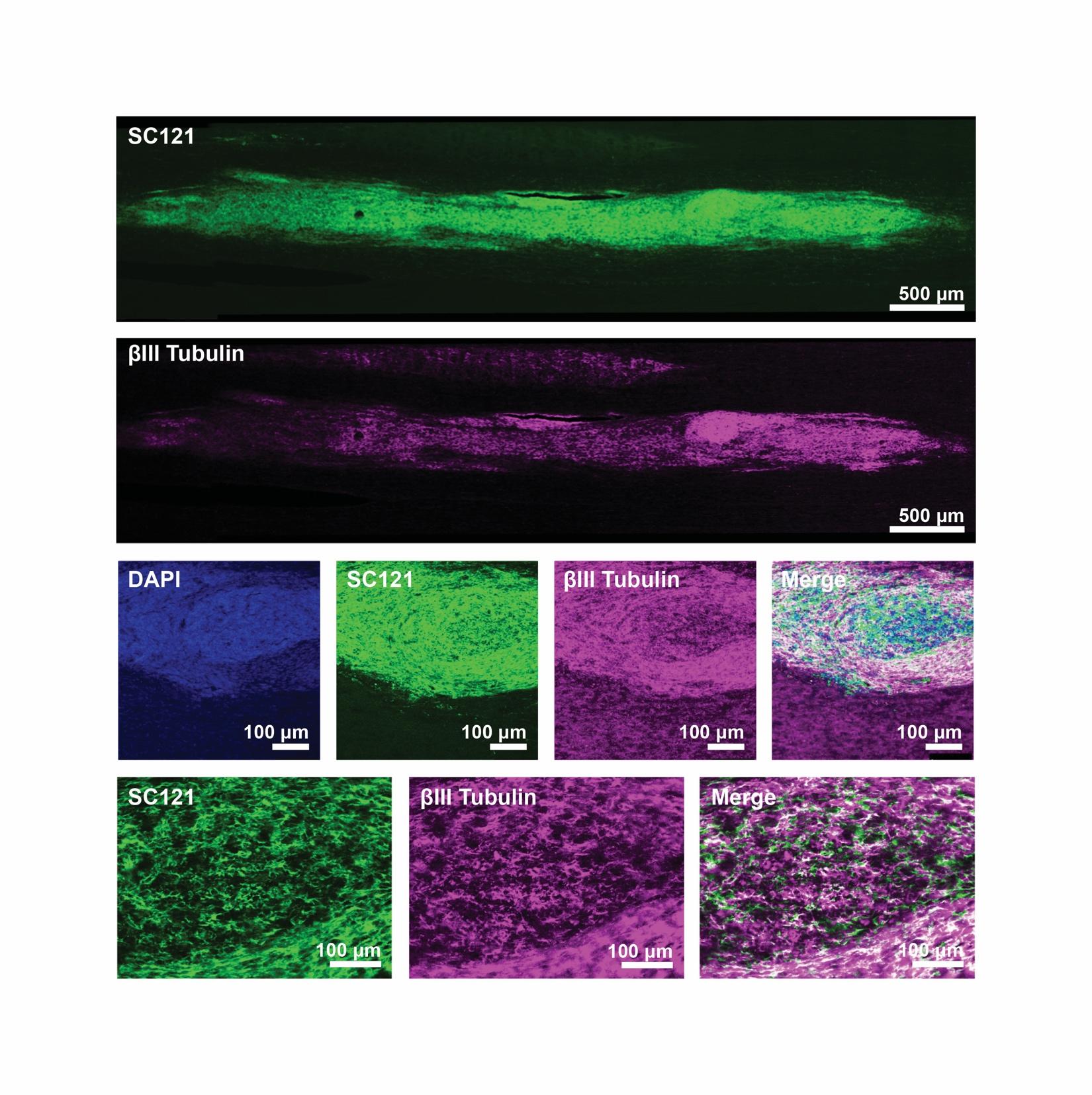

**Figure S6.** *Transplanted hiPSC-DCNs maintain neuronal phenotype.* Representative images taken from the transverse plane in a rat transplanted with hiPSC-DCNs delivered in SHIELD. DAPI is in blue, SC121 (human cytoplasmic marker) is in green, βIII tubulin (neuronal marker) is in purple.
